# The mechanism of DNA sensor cGAS-STING regulating autophagy-related signal pathways

**DOI:** 10.1101/2022.07.05.498918

**Authors:** Wen Song, Meng Sun, Yuchun Liu, Yizhe Zhang, Haiwei Lou, Huimin Fang, Qi Guo

## Abstract

Cyclic GMP-AMP synthase (cGAS) serves as a DNA sensor for recognizing and binding microbial or self-DNA molecules in cells. Upon binding DNA, cGAS produces the second messenger cGAMP to activate the stimulator of interferon genes (STING), which mediates the expression of type 1 interferons (IFN-I) and other cytokines. cGAS-STING-mediated signal pathway plays an important role in innate immune reaction against microbial infections as well as autoimmunity, tumor immunology, and cellular senescence. During this process, cGAS regulates DNA damage repair and induces STING-mediated NF-κB and MAPK signal pathways in autophagy and lysosome-dependent cell apoptosis. However, the molecular mechanisms of cGAS-STING-mediated autophagy still need to be explored. Here, we found that cGAS-STING promotes autophagy by influencing multiple autophagy-related signal pathways: First, cGAS-STING promotes autophagy by inhibiting the mTOR signaling pathway; Second, cGAS-STING affects autophagy by regulating autophagy-related proteins that are also involved in cGAS-STING-mediated IFN-I and NF-κB signaling pathways; Third, the Bcl2 can interact with cGAS and STING to induce cGAS-STING-mediated up-regulation of IFN-I signaling pathway and down-regulation of NF-κB. In addition, we also found that USP19 can significantly reduce the K11-linked ubiquitination of STING in the process of autophagy. Our findings unveiled the functional role and the mechanism of cGAS-STING signaling in regulating autophagy.

## 1. Introduction

In mammals, the first line of defense against foreign genetic material is the innate immune system. The cGAS-STING signaling pathway, discovered in recent years, is crucial for recognizing intracellular double-stranded DNA and inducing natural immune responses[1]. After infections, increased intracellular DNA is detected by the pathway that involves cGAS[2]. In the cGAS-STING signaling pathway, cGAS acts as a receptor for recognizing double-stranded DNA and forms a dimeric cGAS-DNA complex, which catalyzes the formation of the second messenger 2’3’-cGAMP and transmits the signal to the endoplasmic reticulum protein STING[3]. The activated STING signal recruits TBK1 to stimulate the transcription factors NF-kB and IRF3 to activate IFN-I signaling response[4].

Recent research has reported extensive mechanisms of the cGAS-STING pathway mainly mediated by four independent STING functions: IRF3 activation, NF-κB activation, autophagy regulation, and induction of lysosomal cell death (LCD) via STING lysosomal trafficking[1]. More recent data indicate that scientists have explored the relationship between several signaling pathways. Wang et al. proposed that the inflammasome induces caspase 1 to cleave cGAS, and cleaved cGAS cannot recognize DNA, thereby inhibiting type I IFNs signaling responses in the inflammatory body signaling pathway[5]; UNC-51-like kinase 1 (ULK1) has been reported to phosphorylate STING to inhibit the activity of IRF3 and affect IFN-I signaling pathway[6]; Beclin-1 interacts with cGAS and inhibits enzyme activity of cGAS, thereby hindering the production of cGAMP and increasing autophagy[7]. ULK1 and Beclin-1 are important autophagic proteins in the initiation and development of autophagy. Beclin-1 contains a Bcl2 binding domain, a coiled-coil domain, and an evolutionarily conserved domain, which interact with Bcl2, Atg14L, and PI3KC3, respectively[8].

Beclin-1 interacts with the apoptotic molecule B-cell lymphoma 2 (Bcl2), which is suppressed and in a resting state in normal cells [1]. When the autophagy signal is generated, Beclin-1 is separated from Bcl2 to facilitate the initiation of autophagic vacuole formation[9]. Moreover, it was reported that the autophagy protein ATG9a colocalizes with STING and disrupts the aggregation of STING and TBK1 to down-regulate the type I IFN signaling pathway[8, 10]. Compared with wild type macrophages, there was a significant decrease in the colocalization of selective autophagy markers with DNA in cGAS^-/-^ and STING^-/-^ macrophages[11]. It suggested that cGas-STING pathway plays an ancient and highly conserved role in autophagy.

The mechanistic target of rapamycin (mTOR) is an important protein during the initiation of autophagic vacuole formation. When nutrient is rich, the key nutrient sensor mTOR complex 1 (mTORC1) can be activated to suppress Ulk1 activation by phosphorylating Ulk1 Ser 757[4, 12], which disrupts the interaction between Ulk1 and AMPK and inhibits autophagosome assembly[13, 14]. As reported, the cGAS-STING pathway could regulate canonical autophagy through mTOR-Beclin1 pathway and via ER stress-mTOR signaling[10].

Most important molecules in the natural immune response, such as TBK1, IRF3, BECN1, ULK1, et al., could be phosphorylated or ubiquitinated during the autophagy process. Park et al. showed that ULK1 phosphorylates ATG14 to induce a strong autophagy response in the absence of nutrients or mTORC1 inhibition[15]. In addition, AMPK has a dual role in regulating the VPS34–Beclin-1 complex: one is that AMPK can promote the activity of Beclin-1; the other one is that AMPK phosphorylates Beclin-1 to inhibit the activity of VPS34-Beclin-1[16]. It has been confirmed that STING is subsequently phosphorylated by ULK1 caused by an increased IRF3 via NF-κB signaling[6], which plays an important role in STING signaling pathways. Ubiquitin has recognized as an important regulatory mechanism for cellular signaling. It is found that lysine 48 (K48)-linked proteins are destined for proteasomal degradation, while K11, K27, or K63-linked proteins represent post-translational modifications. Nazio et al. mentioned that TRAF6 facilitates ULK1 ubiquitination, which promotes oligomerization of ULK1 to enhance autophagy under the assistance of Ambra1 (activating molecule in Beclin1-regulated autophagy)[17]. The TRIM14-USP14 molecular complexes inhibit P62-mediated degradation of cGAS by inhibiting cGAS K48-linked ubiquitination to positively regulate the antiviral immunity of DNA viruses[18]. TRIM56 interacts with STING to produce K63-linked ubiquitination, which promotes dimerization of STING, and recruits TBK1, causing the production of downstream IFN-β[19]. Above all, phosphorylation and ubiquitination are both the main focus of current research.

In this study, we found that cGAS-STING could induce autophagy through the mTOR signaling pathway in 293t and HeLa cells. Then, we found that autophagy-related protein Bcl2 interacts with Beclin-1, which results in down-regulation of the cGAS-STING-mediated IFN I signaling pathway. In addition, we also found that USP19 could decrease K11-linked ubiquitination of STING in the process of autophagy. Our findings unveiled the functional role and the regulatory mechanism of cGAS-STING signaling in autophagy.

## 2. Materials and Methods

### 2.1 Constructs

pENTRY, pcDNA3.1, NF-κB and ISRE promoter luciferase reporter plasmids, mammalian expression plasmids for HA- or Flag-tagged RIG-I (2CARD), cGAS, STING, ATGs, and DUBs were provided by Jun Cui (Sun Yat-sen University).

### 2.2 Cell culture and reagents

HEK293T (human embryonic kidney 293T), HeLa cells were cultured in DMEM (Gibco) with 10% (vol/vol) fetal bovine serum (Gibco). The experiment of cell culture was performed as described[20].

### 2.3 Antibodies

The following antibodies were used in this study: anti-IRF3 (sc-9082), goat anti-rabbit IgG-HRP (sc-2004), goat anti-mouse IgG-HRP (sc-2005) (Santa Cruz Biotechnology); HRP-anti-Flag (M2) (A8592) and anti-β-actin (A1978) (Sigma); HRP-anti-hemagglutinin (clone 3F10), anti-Myc-HRP (11814150001)(Roche Applied Science); anti-LC3B(#3868), anti-p62(#39749), anti-TBK1 phosphorylated at Ser172 (#5483), anti-p70S6K phosphorylated at Ser371 (#9208), anti-p70S6K(#2708), anti-ULK1(#8054), anti-ULK1 phosphorylated at Ser757 (#14202) (Cell Signaling Technology).

### 2.4 Immunoprecipitation and immunoblot analysis

For immunoprecipitation, whole cell extracts were prepared after transfection or stimulation with appropriate ligands, followed by incubation overnight with the appropriate antibodies plus Protein A/G beads (Pierce). And anti-Flag, anti-HA beads, anti-Myc beads (Sigma) for immunoprecipitation. Beads were washed five times with low-salt lysis buffer, and immunoprecipitants were eluted with 3× SDS Loading Buffer and resolved by SDS-PAGE. Proteins were transferred to PVDF membranes (Bio-Rad) and further incubated with the appropriate antibodies. Immobilon Western Chemiluminescent HRP Substrate (Millipore) was used for protein detection.

### 2.5 Luciferase and reporter assays

HEK293T (2 × 105) cells were plated in 24-well plates and transfected with plasmids encoding an ISRE or NF-κB luciferase reporter (firefly luciferase; 100 ng) and pRL-TK (renilla luciferase plasmid; 10 ng) together with 100 ng plasmid encoding Myc-cGAS, and mutants of Myc-cGAS(NT), Myc-cGAS(CT), Myc-cGAS(211/213), Myc-cGAS(174), Myc-cGAS(384), Myc-cGAS(394), Myc-cGAS(407), Myc-cGAS(411), HA-STING, and HA-STING 1-340, Flag-Bcl2 and increasing concentrations (0, 10, 20, 50, 100, or 200 ng) of plasmids. Empty pcDNA3.1 vector was used to maintain equal amounts of DNA among wells. Cells were collected at 24 h after transfection and luciferase activity was measured with the Dual-Luciferase Assay (Promega) with a Luminoskan Ascent luminometer (Thermo Scientific) according to the manufacturer’s protocol. Reporter gene activity was determined by normalization of the firefly luciferase activity to renilla luciferase activity.

### 2.6 Statistical analysis

Data are represented as mean ± SD when indicated, and Student’s t-test was used for all statistical analyses with the GraphPad Prism 5.0 software. Differences between groups were considered significant when P-value was < 0.05.

## 3. Results

### 3.1 cGAS-STING induces the autophagy independently with the type I IFN signaling pathway

To explore whether cGAS or STING independently induces autophagy, we overexpressed cGAS and STING in HEK393T or Hela cells. Detecting autophagy makers of LC3B and p62, we found that STING cannot directly trigger the occurrence of autophagy in 293T cells, but it induces the autophagy after the coexpression of cGAS (Fig. 1A). Conversely, cGAS cannot directly stimulate autophagosome formation unless there is a coexpression of STING (Fig. 1B). In addition, autophagy was significantly enhanced by the co-overexpression of cGAS and STING in Hela cells (Fig. S1A).

**Figure 1.**
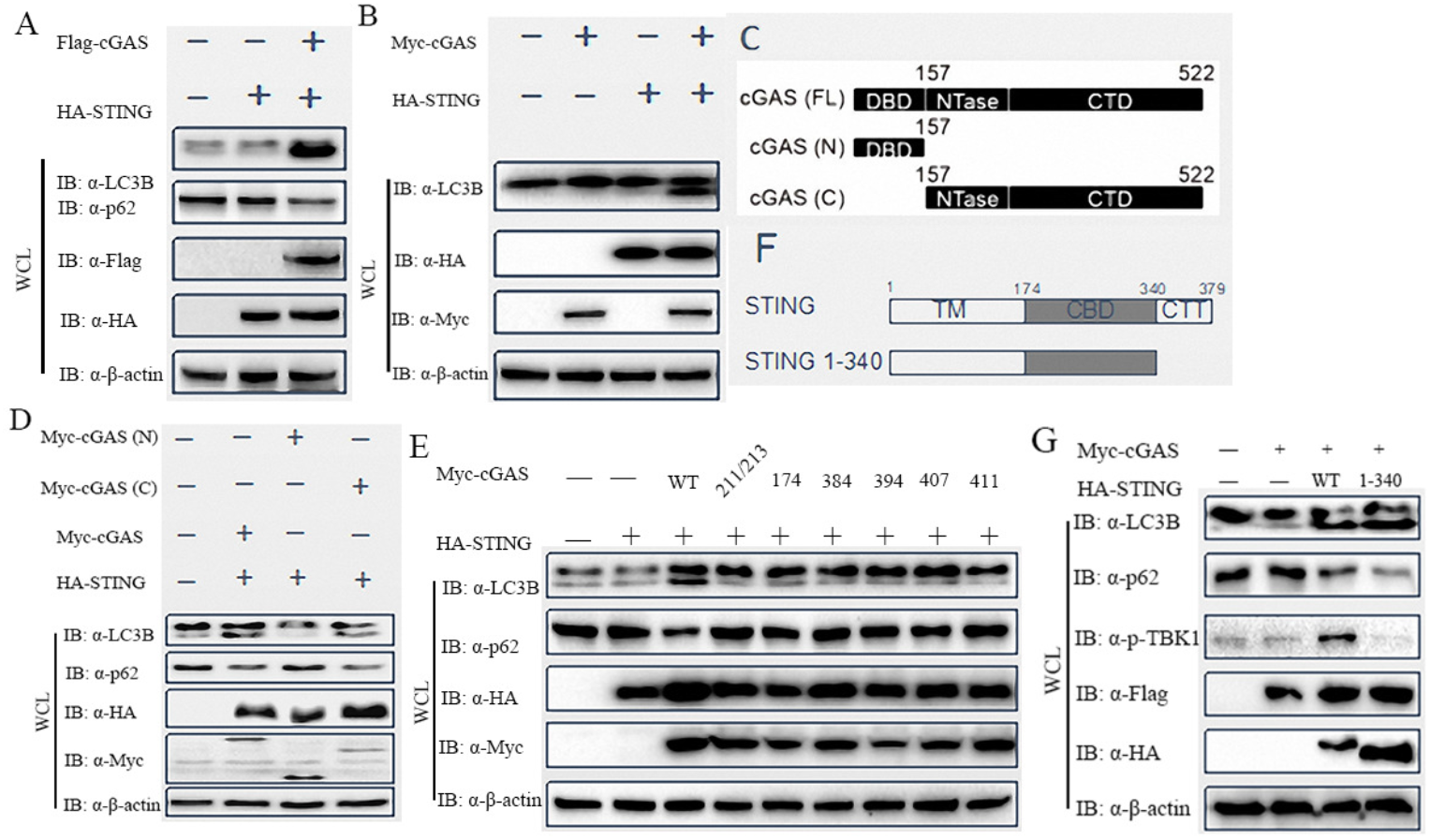
cGAS and STING work together to induce autophagy. **(A)** Immunoblot of autophagy makers LC3B and p62 after overexpressing HA-STING, or HA-STING and Flag-cGAS. **(B)** Immunoblot of autophagy makers LC3B after overexpressing HA-STING, Myc-cGAS, or Flag-cGAS and HA-STING. **(C)**: Diagram of cGAS domain structures and truncation constructs of cGAS subdomains (FL, N, or C). **(D)** Immunoblot of autophagy makers LC3B and p62 after co-overexpressing HA-STING and My-cGAS or N- and C-truncation constructs. **(E)** Immunoblot of autophagy makers LC3B and p62 after co-overexpressing HA-STING (WT) and Myc-cGAS carrying mutations of residues 211/213,174, 384, 394, 407, or 411, which are important for its enzymatic activity. **(F)** Diagram of STING domain structures and its C-terminal truncation construct. **(G)** Immunoblot of p-TBK1 and autophagy makers LC3B and p62 after co-overexpressing Myc-cGAS (WT) and HA-STING constructs.

It was reported that cGAS-STING activates downstream molecules of the IFN-I pathway depending on the functional regions of the cGAS and its enzymatic activity[21]. To investigate whether different cGAS regions have a certain function during autophagy, we generated the cGAS truncations as shown in Fig.1C. Then, we transfected different cGAS truncation constructs and STING into 293T cells. We found that the functional domains of cGAS induces autophagy and plays a critical role in activating downstream molecules of the IFN-I signaling pathway (Fig. 1D, Fig. S1B). Furthermore, we studied all mutation sites affecting cGAS enzyme activity and found that none of them causes obvious autophagy (Fig. 1E). Moreover, these mutants of cGAS cannot activate of the type I IFN downstream signaling pathway (Fig. S1C).

The C-terminal domain of STING contains dimerization regions and the carboxyl-terminus, which is a functional region to activate the downstream signal pathways. To explore the certain function of STING functional regions, we divided the STING into sections as shown in Fig. 1F. Then we overexpressed those sections in HEK293T cells to observe the autophagy-related markers. The results suggested that the N-terminus of STING causes stimulation of autophagy (Fig. 1G), as well as activation of the IFN-I signaling pathway (Fig. S1D). Activated STING induces TBK1 to phosphorylate itself, STING, and IRF3 in IFN-I signaling response. According to the results in this study, deletion of C-terminus in STING 1-340 did not activate the phosphorylation of TBK1 or increased the expression of type I IFN but caused enhanced autophagy (Fig. 1G). Therefore, we speculated that the mechanisms by which cGAS-STING promotes autophagy and the IFN-I signaling pathway are independent of each other.

### 3.2 cGAS-STING regulates autophagy by inhibiting mTOR signaling pathway

mTOR signaling pathway plays roles in cGAS-STING mediated autophagy initiation as well as IFN-I signaling response in the innate immune system. Are there cross-talks between mTOR and cGAS-STING signaling pathways in the process of autophagy? To address this question, we first studied the expression of phosphorylated P70S6K and ULK1, which are downstream molecules in the mTOR signaling pathway. We found in the case of co-expressing cGAS and STING, the phosphorylation of the downstream molecules p70S6K Thr389 and ULK1 Ser757 in the mTOR signaling pathway was significantly inhibited (Fig. 2A). Furthermore, we also found that the phosphorylation inhibition was still obvious after the addition of RAPA, the mTOR signaling pathway inhibitor (Fig. 2B). Then, we analyzed the change in p70S6K Thr389 and ULK1 Ser757 phosphorylation after two different STING regions were overexpressed in 293T, and found that the C-terminal truncation of STING could also reduce phosphorylation of downstream molecules of mTOR (Fig. 2C). As such, cGAS-STING mediated autophagy by affecting the phosphorylation of the downstream molecules P70S6K and ULK1 in the mTOR signaling pathway, which was dependent on STING.

**Figure 2.**
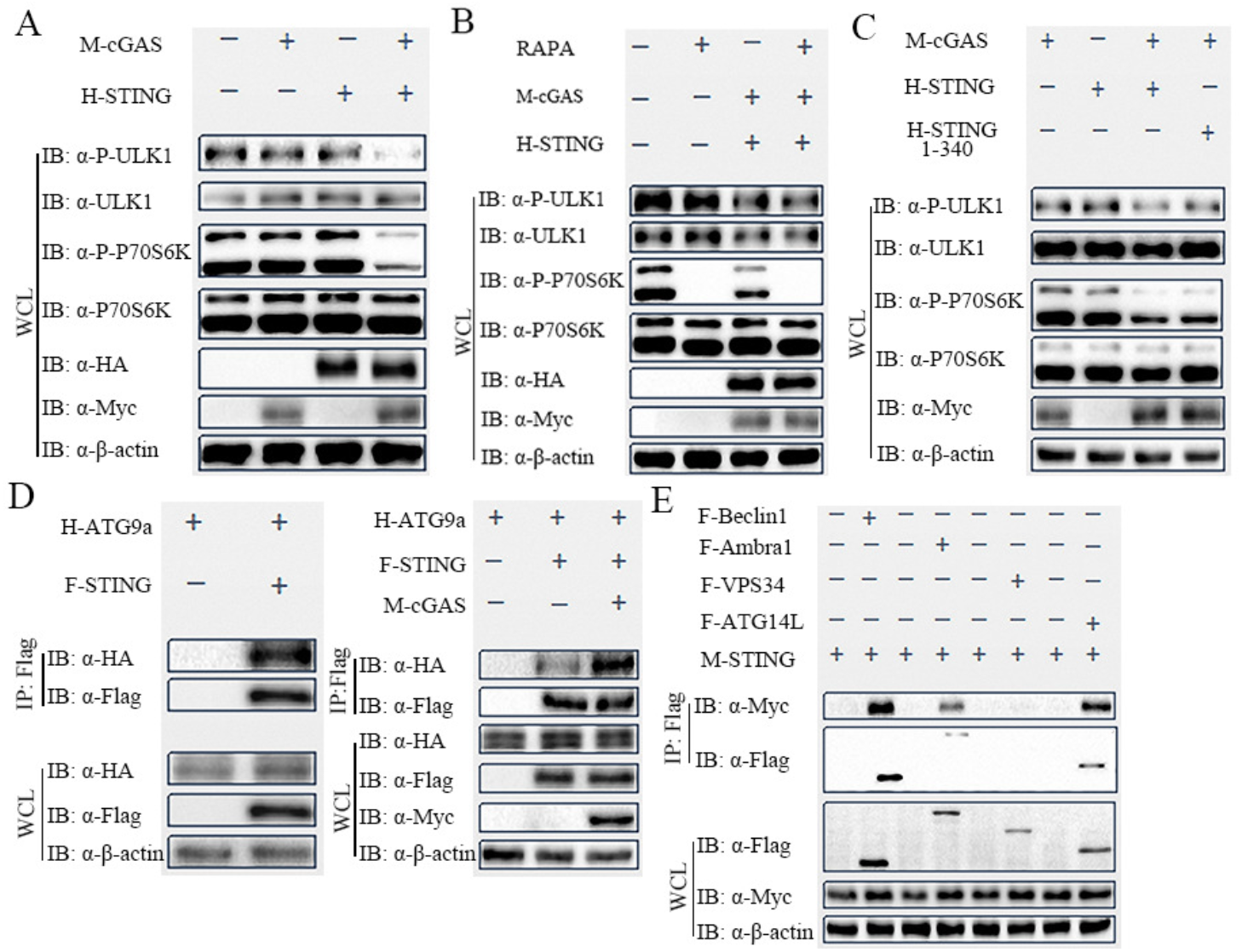
The effect of cGAS-STING on autophagy-related proteins. **(A)** Immunoblot of autophagy-related proteins in the mTOR signaling pathway after overexpressing Myc-cGAS, or HA-STING, or both proteins. **(B)** Immunoblot of autophagy-related proteins in the mTOR signaling pathway in cGAS and STING overexpressing HEK293T cells without or with rapamycin (RAPA, 100 nM) treatment for 24 h. **(C)** Immunoblot of autophagy-related proteins in the mTOR signaling pathway in cGAS and STING or the truncated STING (1-340 aa) overexpressing cells. **(D)** Analyzing of interaction between HA-ATG9a and Flag-STING by co-IP with anti-Flag beads and immunoblot with anti-HA after overexpressing HA-ATG9a and Flag-STING (left panel) or with the coexpression of Myc-cGAS (right panel). **(E)** Analyzing the interaction of STING with the Beclin-1 complex members (Flag-ATG14L, Flag-Beclin-1, Flag-VPS34, Flag-Ambra1).

### 3.3 Bcl2 regulates cGAS-STING-mediated type I IFN signaling

In addition, we also conducted studies on STING and proteins in the autophagy signaling pathway and found the interaction between STING and various autophagy proteins. We found that STING interacted with ATG9a, Beclin-1, ATG14L, and Ambra1 in the Beclin-1 complex (Fig. 2D&E). cGAS could enhance the interaction between ATG9a and STING (Fig. 2D). Beclin-1 interacts with Bcl2 in the initiation of autophagy, and we found that there is an interaction between Beclin-1 and STING (Fig. 2E, Fig. S2A). Based on the interaction between Bcl-2 and Beclin-1, we have developed a strong interest in the relationship between Bcl-2 and cGAS and the role of Bcl-2 on the IFN-I signaling pathway. We over-expressed Bcl2 and cGAS-STING in 293T cells and found there was indeed an interaction between Bcl2 and cGAS/STING (Fig. 3A&B). This conclusion was also confirmed in the Hela cell line (Fig. 3B and S2B).

**Figure 3.**
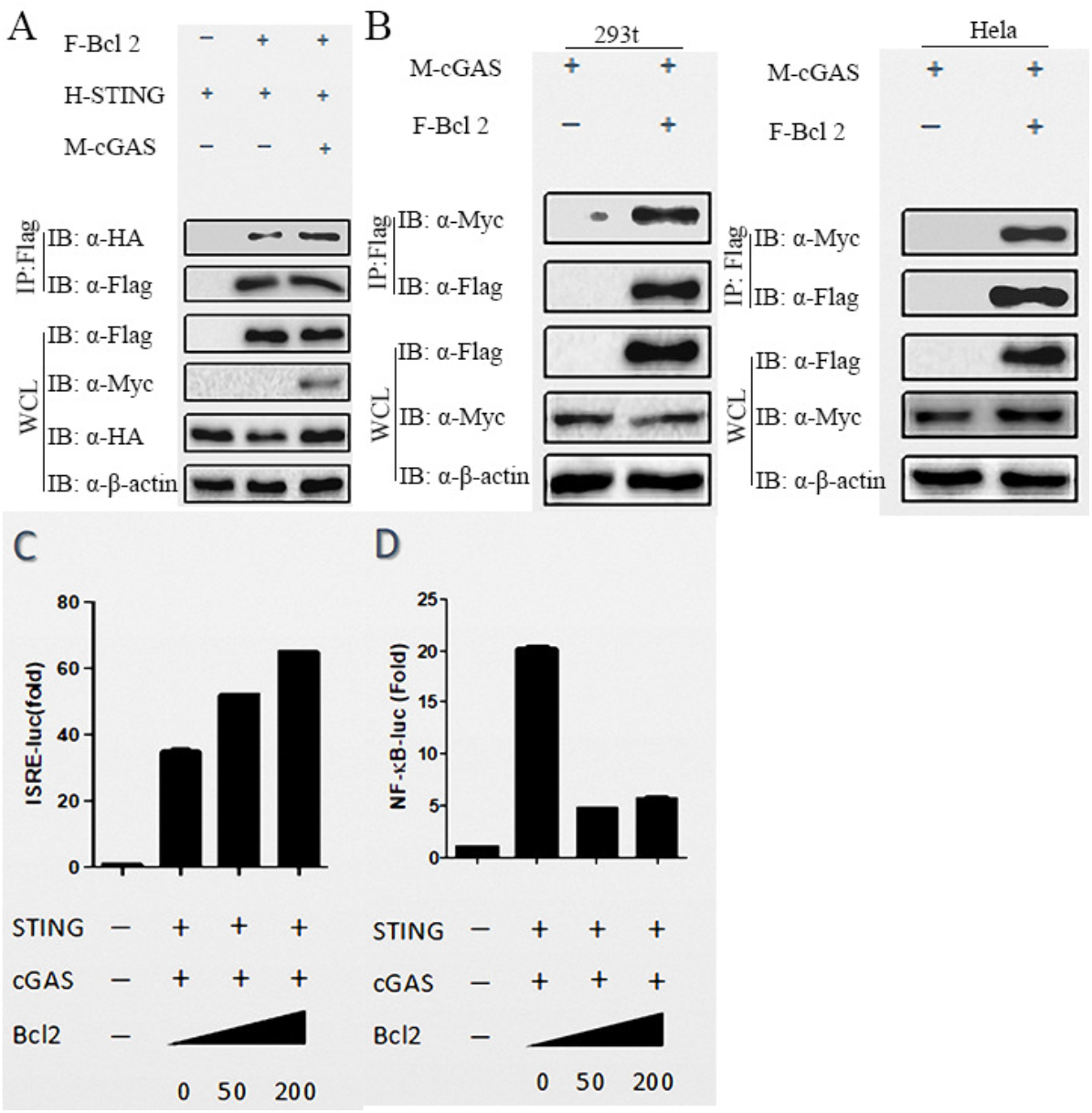
Bcl2 affects cGAS/STING-mediated type I IFN signaling. **(A)** Interaction analyses of HA-STNG and Flag-Bcl2 with or without overexpressing Myc-cGAS. **(B)** Interaction analyses of Myc-cGAS and Flag-Bcl2 in HEK293T or Hela cells. (**C** and **D**) Luciferase activity analyses in HEK293T cells transfected with a ISRE luciferase (ISRE-luc) reporter or a NF-κB luciferase (NF-κB-luc) reporter construct with the control plasmid, Myc-cGAS, HA-STING, and an increasing amount of FLAG-Bcl2 expression plasmids.

To explore whether Bcl2 induces IFN-I or NF-κB signaling pathways through cGAS-STING, we performed experiments on the relationship between Bcl2 and cGAS-STING-mediated IFN-I signaling in 293T cells. The results showed that after the addition of Bcl2, the IFN-I signaling in cells was significantly up-regulated, while the NF-κB pathway was down-regulated (Fig.3C&D). Therefore, we concluded that Bcl2 interacts with STING as well as with cGAS to induce cGAS-STING-mediated up-regulation of the IFN-I signaling pathway.

### 3.4 The ubiquitination of STING in cGAS-STING mediated autophagy

Next, we studied ubiquitination in the cGAS-STING signaling pathway to explore its relationship with autophagy induction. We found the ubiquitination of STING, especially K11-linked ubiquitination of STING was significantly reduced while cGAS and STING were overexpressed (Fig.4A&B). USP19 stabilizes Beclin-1 by removing the K11 ubiquitin chain of Beclin-1 to promote the formation of the Beclin-1 complex, thereby altering autophagy formation[22]. Then, we investigated if there was a function of USP19 on the ubiquitination of STING. We found that USP19 interacts with STING in autophagy (Fig.4C) and inhibits the ubiquitination of STING, especially K11-linked ubiquitination of STING (Fig.4D).

**Figure 4.**
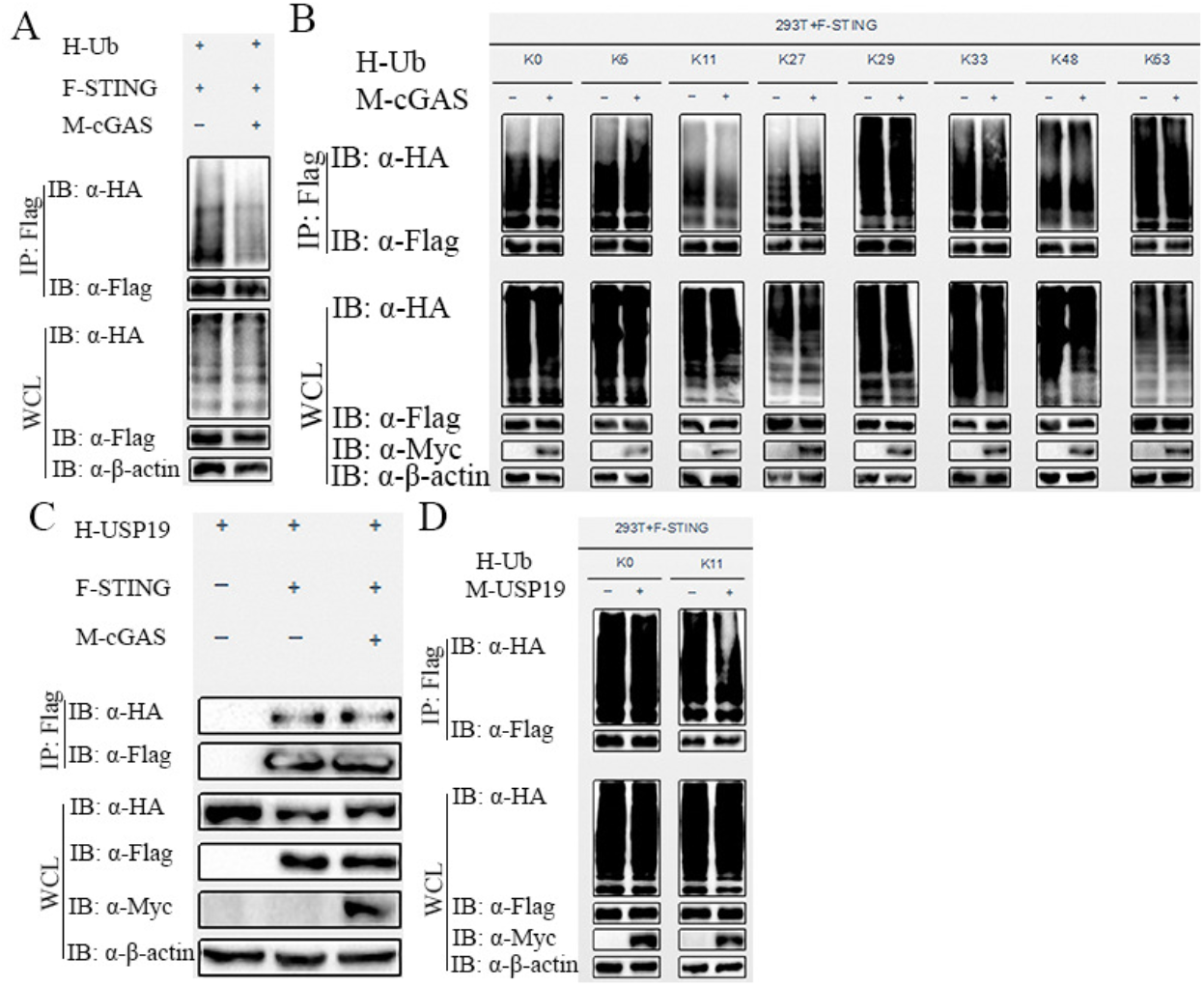
The ubiquitination of STING in cGAS-STING mediated autophagy. (**A and B**) Analyzing the ubiquitination of STING after overexpressing Flag-STING and HA-ubiquitin (**A**, WT) and its mutant variants (**B**: K0, K6, K11, K27, K29, K33, K48, K63-linked ubiquitin) without or with the coexpression of Myc-cGAS. **(C)**Analyzing the interaction between HA-USP19 and Flag-STING by co-IP and immunoblot without or with the coexpression of Myc-cGAS. **(D)** Analyzing the ubiquitination of STING after overexpressing Flag-STING by HA-Ub K0 or HA-Ub K11 without or with Myc-USP19 coexpression.

## 4. Discussion

Involved in a variety of autophagy-related proteins, autophagy is the internal nature of the normal process in dealing with the cytoplasm of the nonfunctional components in the initial stage[23]. Autophagy is also crucial for the innate immune system to remove exotic or intrinsic substances. cGAS/STING is a receptor that recognizes DNA, and plays an important role in the innate immune system: cGAS recognizes DNA and produces cGAMP to transmit the signal to STING, then activates TBK1 or IRF3 to induce type I IFN or NF-κB signaling pathways[4, 24]. Kato and Burdette explored the cGAS functional domain and found the enzyme sites and domains of cGAS[21, 25]. Therefore, we focused on the functional domains of cGAS and STING in the study of the relationship between cGAS-STING signaling and autophagy. We found the activation of cGAS-STING depends on STING 1-340 domain and C-terminal regions of cGAS. Besides, we studied that the functional domains of cGAS-STING, which are important for inducing autophagy.

Then we focused on the relationship between cGAS/STING and mTOR signaling pathway. mTORC1 is an important key mediator in the formation of autophagosomes. Inhibition of mTORC1 can promote the formation of the ULK1 complex. With the activation of ULK1 complex and Beclin-1 complex, autophagosome undergoes initiation, extension, and formation to eventually fusion with lysosomes [17]. Therefore, we investigated the role of mTOR signal in cGAS-STING-mediated autophagy. We found that cGAS-STING signal promotes autophagy by regulating the phosphorylation of ULK1 and P70S6K, which are downstream of mTOR signaling pathway.

We also explored the interaction between cGAS-STING and some autophagy-related molecules in the autophagic signaling pathway. ULK1 was reported to phosphorylate STING to inhibit the activity of IRF3 and induce type I IFN signaling pathway[6]; Beclin-1 can inhibit the cGAS to decrease the production of cGAMP and IFN-β, while ATG9a blocks the aggregation of STING and TBK1 through co-localization with STING[26]. Furthermore, we found that STING could be targeted on multiple autophagy-related proteins, such as the Beclin1 complex and ATG9a. In addition, we also found that STING and Beclin-1 had an inseparable interaction, while Beclin-1 and Bcl-2 interact with each other. The Bcl-2 inhibits the signaling transmission of Beclin-1 to the downstream of autophagy. Bcl2 could also up-regulate cGAS-STING-mediated type I IFN signaling pathway and down-regulate cGAS-STING-mediated NF-κB signaling pathway, which may be due to the relationship between Bcl2 and Beclin-1.

As known, cGAS-STING promoted autophagy via the mTOR signaling pathway or autophagy-related proteins, but how is cGAS-STING modified while interacting with molecules and transmitting downstream signals? During the process of autophagy, phosphorylates Beclin-1 S91/94, as well as Beclin-1 S14 phosphorylation by ULK1 plays an important role[6]. The lysine at position 150 of STING undergoes K11-linked ubiquitination in the presence of RNF26 to limit strong type I IFN response[27]. Therefore, we investigated the ubiquitination of STING in cGAS-STING-mediated autophagy and found that the ubiquitination of STING was significantly reduced, especially K11-linked ubiquitin after the overexpression of cGAS and STING. We also found that USP19 inhibits the K11-linked ubiquitination of STING, which was consistent with the previous findings[22]. However, there still needs more evidence to confirm whether K11-linked ubiquitinated by USP19 is important for autophagy.

Autophagy closely relates to allogeneic phagocytosis, nutritional deficiencies, infections, cell repair mechanisms, and programmed death. Autophagy not only maintains homeostasis by degrading non-essential substances for energy or eliminating damaged molecules in vivo, but also plays an important role in a variety of diseases, such as osteoarthritis, cancer, and Parkinson’s disease et al. Therefore, the further investigation focused on mechanisms of cGAS-STING signaling and autophagy could be useful for understanding the natural immunity and autophagy and provide the theoretical basis for subsequent immunotherapy or drug treatment.

## Acknowledgments

This work was supported by Bingtuan Science and Technology Project [2019AB034], Scientific and Technological Innovation Leaders in Central Plains of Henan (194200510002).

## Author Contributions

Qi Guo and Huimin Fang designed the study; Yizhe Zhang and Haiwei Lou interpreted data; Wen Song and Meng Sun wrote the manuscript; Wen Song, Meng Sun and Yuchun Liu performed experiments and analyzed data.

